# BITACORA: A comprehensive tool for the identification and annotation of gene families in genome assemblies

**DOI:** 10.1101/593889

**Authors:** Joel Vizueta, Alejandro Sánchez-Gracia, Julio Rozas

## Abstract

Gene annotation is a critical bottleneck in genomic research, especially for the comprehensive study of very large gene families in the genomes of non-model organisms. Despite the recent progress in automatic methods, the tools developed for this task often produce inaccurate annotations, such as fused, chimeric, partial or even completely absent gene models for many family copies, which require considerable extra efforts to be amended. Here we present BITACORA, a bioinformatics solution that integrates sequence similarity search tools and Perl scripts to facilitate both the curation of these inaccurate annotations and the identification of previously undetected gene family copies directly from DNA sequences. We tested the performance of the BITACORA pipeline in annotating the members of two chemosensory gene families of different sizes in seven available chelicerate genome drafts. Despite the relatively high fragmentation of some of these drafts, BITACORA was able to improve the annotation of many members of these families and detected thousands of new chemoreceptors encoded in genome sequences. The program generates an output file in the general feature format (GFF) files, with both curated and novel gene models, and a FASTA file with the predicted proteins. These outputs can be easily integrated in genomic annotation editors, greatly facilitating subsequent manual annotation and downstream evolutionary analyses.

## Introduction

The falling cost of high-throughput sequencing (HTS) technologies made them accessible to small labs, promoting a large number of genome-sequencing projects even in non-model organisms. Nevertheless, genome assembly and annotation, especially in eukaryotic genomes, still represent major limitations (Dominguez Del Angel et al., 2018). The unique genomic characteristics of many non-model organisms, often lacking pre-existing gene models (Yandell & Ence, 2012), and the absence of closely related species with well-annotated genomes, converts the annotation process in a big challenge. The state-of-the-art pipelines for *de novo* genome annotation, like BRAKER1 or MAKER2, allow integrating multiple evidences, such as RNA-seq, EST data or gene models from other annotated species (using for example GeneMark, Exonerate, or GenomeThreader) with *ab initio* gene predictions (from Augustus or SNAP) in order to produce structural annotations of genome sequences (Gremme, Brendel, Sparks, & Kurtz, 2005; Hoff et al., 2016; Holt & Yandell, 2011; Korf, 2004; Lomsadze, Burns, & Borodovsky, 2014; Slater & Birney, 2005; M. Stanke & Waack, 2003; Mario Stanke, Diekhans, Baertsch, & Haussler, 2008). Some of these pipelines, such as BRAKER1, will only report those gene models with evidences. However, the gene models predicted by these automatic tools are often inaccurate, especially those belonging to gene families. Their curation frequently requires the use of additional programs, such as Augustus-PPX (Keller, Kollmar, Stanke, & Waack, 2011), or semi-automatic approaches evaluating the quality of supporting data. This latter task is usually performed in genomic annotation editors, such as Apollo, which give researchers the option to work simultaneously in the same annotation project (Lee et al., 2013).

There are a number of issues affecting the quality of gene family annotations, especially for either old or fast evolving families (Yohe et al., 2019). First, new duplicates within a family usually originate by unequal crossing-over and are found in tandem arrays in the genome, being the more recent duplicates also the physically closest (Clifton et al., 2017; Vieira, Sánchez-Gracia, & Rozas, 2007). This configuration often causes local miss-assemblies that result in the incorrect or failed identification of tandem duplicated copies (i.e., it produces artifact, incomplete, or chimeric genes along a genomic region). Secondly, the identification and characterization of gene copies in medium- to large- sized families tends to be laborious, requiring data from multiple sources, including well-annotated remote homologs and hidden Markov model (HMM) profiles. Certainly, the fine and robust identification and annotation of the complete repertory of a gene family in a typical genome draft is a challenging task that requires important additional efforts, which are very tedious to perform manually.

In order to facilitate this curation task, we have developed BITACORA, a bioinformatics pipeline to assist the comprehensive annotation of gene families in genome assemblies. BITACORA requires of a structurally annotated genome (GFF and FASTA format), and a curated database with well-annotated members of the focal gene families. The program will perform comprehensive BLAST and HMMER searches (Altschul, 1997; Eddy, 2011) to identify putative candidate gene regions (already annotated, or not), combine evidences from all searches and generate new gene models. The outcome of the pipeline consists of a new structural annotation (GFF) file along with their encoded sequences. These output sequences can be directly used to conduct downstream functional or evolutionary analyses, to be included as evidences in other annotation pipelines (BRAKER1 or MAKER2; Hoff et al., 2016; Holt & Yandell, 2011), to improve existing gene models predictions, or to facilitate a fine re-annotation in genome browsers such as Apollo (Lee et al., 2013).

## Methods and implementation

### Input data files

BITACORA requires: i) a data file with the genome sequences (in FASTA format), ii) the associated GFF file with annotated features (either in GFF3 or GTF formats; features must include both transcript or mRNA and CDS), iii) a data file with the predicted proteins included in the GFF (in FASTA format), and iv) a database (here referred as FPDB database) with the protein sequences of well annotated members of the gene family of interest (focal family; in FASTA format) along with its HMM profile (see Supplementary Material for a detailed description of FPDB construction). Since sequence similarity-based searches are very sensitive to the quality of the proteins in FPDB, it is important to include in this database highly curated proteins from closely related species. This is especially important for the annotation of very old or fast-evolving gene families. Also, the use of a HMM profile increases the likelihood of identifying sequences encoding new members; these profiles can be obtained from external databases (such as PFAM) or build using high quality protein alignments with the program *hmmbuild* (Finn *et al.*, 2014). Before starting the analysis, BITACORA checks whether input data files are correctly formatted; otherwise, it will suggest some format converters distributed with the program (see Troubleshooting section in Supplementary Material).

### Curating existing annotations

The BITACORA workflow is divided in three main steps (Fig. 1). The first step consists in the identification of all putative homologs of the FPDB sequences from the focal gene family that are already present in the input GFF file, and the curation of their gene models (referred hereinafter as b-curated (bitacora-curated) gene models or proteins). Specifically, the pipeline launches BLASTP and HMMER searches (Altschul, 1997; Eddy, 2011) against the proteins predicted from the features in the input GFF using the FPDB protein sequences and HMM profiles as queries; the resulted alignments are filtered for quality (i.e. BLASTP hits covering at least two-thirds of the length of query sequences or including at least the 80% of the complete protein used as a subject are retained). The results from both searches are combined into a single integrated result for every single protein (gene model). Then, BITACORA trims the original models based in these combined results, and reports new gene coordinates (b-curated models) in a new updated GFF (uGFF), fixing for example all chimeric annotations. Besides, the proteins encoded by these b-curated models are incorporated to the FPDB (updated FPDB or uFPDB), to be used in an additional search round.

**Fig. 1.**
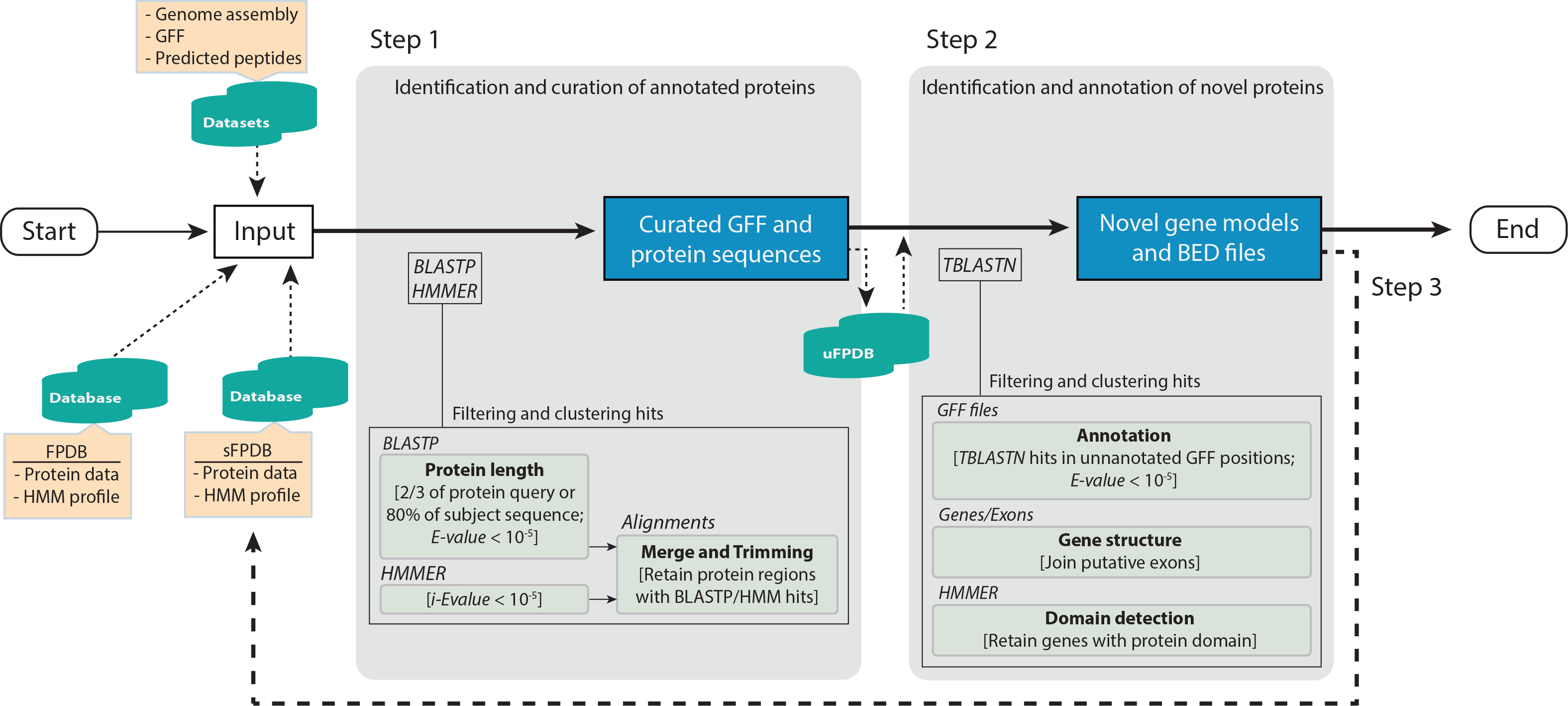
BITACORA workflow.

### Identifying genomic regions encoding new family members

In the second step, BITACORA uses TBLASTN to search the genome sequences for regions encoding homologs of the proteins included in the uFPDB but not annotated in the uGFF. Overlapping TBLASTN hits, which we would expect to represent a unique exon sequence, are merged into one single alignment. Then, all alignments located in the same scaffold and separated less than the maximum allowed intron distance (indicated by the “intron distance parameter”) are connected to obtain a putative single protein coding region (referred hereinafter as b-novel gene models). This step is intended to join coding exons of the same gene based on expected intron distance in the surveyed genome. We provide some scripts to estimate the “intron distance parameter” from the input GFF (see Supplementary Material). Last, to avoid reporting inaccurate b-novel gene models and to identify putative gene fusions among them, BITACORA checks the encoded proteins for the presence of the gene family-specific domain (using the HMM profile in FPDB), and only models having this domain are reported in the final dataset of annotated proteins, tagging those cases that could be the result of a fusion of multiple genes with the label ‘Ndom’ (being N =>2, denoting the presence of more than one protein family domain in the sequence; see Supplementary Material for more details).

### Optional search round and final output

Finally, BITACORA can also be used to perform a second search round using as the input data all proteins obtained in steps 1 and 2 (sFPDB database). This additional step is especially useful for searching remote homologs undetected in the previous steps. The final BITACORA outcome will include therefore, 1) an updated GFF file with both b-curated and b-novel gene models, 2) all non-redundant proteins predicted from these feature annotations (in a FASTA file), 3) two BED files, one with all gene coordinates in the genome sequence and the other with only those regions that encode the novel members of the focal family identified by the program and, 4) all protein sequences found in all steps.

### Additional features

BITACORA could be also used in the absence of either a reference genome for the target species (e.g. for transcriptomic studies) or a precompiled GFF (e.g. for non-annotated genomes); in these cases, the input should be a FASTA file with the set of predicted proteins or the genome sequences, respectively (see Supplementary Material for alternative usage modes). With BITACORA, we also distribute a series of scripts to perform some useful tasks, such as estimating intron length statistics from a GFF, converting GFF to GTF format, and retrieving all protein sequences encoded by the features of a GFF file. Furthermore, to better adjust to the particularities of each genome, BITACORA allows the user to specify the values of most important parameters, such as the *E*-value for BLAST and HMMER searches, the number of threads in BLAST runs, or the maximum intron length required to connect putative exons of the same gene.

### BITACORA application example

As a demonstration of the performance of BITACORA in a group of genomes of different quality and assembly contiguity, we present the extended results of the annotation of two arthropod chemosensory gene families, the insect gustatory receptor (GR) and the Niemann-Pick type C2 (NPC2) gene families (Pelosi et al., 2014; Robertson, 2015), in a subset of seven chelicerate genomes from those analyzed in Vizueta et al., (2018). For the analysis, we retrieved the data (genome sequences, annotations and predicted peptides) of the scorpions *Centruroides sculpturatus* (bark scorpion, genome assembly version v1.0, annotation version v0.5.3; Human Genome Sequencing Center (HGSC)) and *Mesobuthus martensii* (v1.0, Scientific Data Sharing Platform Bioinformation (SDSPB)) (Cao et al., 2013); and of the spiders *Acanthoscurria geniculata* (tarantula, v1, NCBI Assembly, BGI) (Sanggaard et al., 2014), *Stegodyphus mimosarum* (African social velvet spider, v1, NCBI Assembly, BGI) (Sanggaard et al., 2014), *Latrodectus hesperus* (western black widow, v1.0, HGSC), *Parasteatoda tepidariorum* (common house spider, v1.0 Augustus 3, SpiderWeb and HGSC) (Schwager et al., 2017) and *Loxosceles reclusa* (brown recluse, v1.0, HGSC). The GR and NPC2 families show very different protein and genomic features. The GR gene family encodes seven-transmembrane receptors of ~400 amino acids long with an average of 2.3 exons per gene in the genome of the spider *Parasteatoda tepidariorum*; the NPC2 proteins are ~150 amino acids long and have an average of 2.6 exons per gene in the same species.

Strikingly, BITACORA uncovered the identification of thousands of new gene models previously undetected in these chelicerate genomes. For instance, BITACORA was able to identify and annotate 1,234 GR encoding sequences in the bark scorpion *Centruroides sculpturatus*, where only 24 proteins were initially identified by the automatic annotation pipelines (Table 1). Globally, BITACORA identified, annotated and curated 3,371 sequences encoding GR proteins in the seven genomes (3,265 of them absent in structural annotations included in the GFF of these genomes). It is largely known that this gene family evolves rapidly in arthropods, both in terms of sequence change and repertory size, encoding in the same genome very recent and distantly related receptors as well as pseudogenes. Since some of these receptors show a very restricted gene expression pattern (expressed in specialized cells and tissues involved in chemoreception), their transcripts are often missing in RNA-seq data sets, which are one of evidences used for the automatic annotation of the genomes (Joseph & Carlson, 2015; Robertson, 2015; Vizueta et al., 2017; Zhang, Zheng, Li, & Fan, 2014). This fact, added to the huge divergence accumulated between many copies (a mixture of age and rapid evolution), probably prevented the automatic annotation of the GRs uncovered by BITACORA.

**Table 1.** Summary of the number of GRs and NPC2 genes identified by BITACORA in seven chelicerate genomes.

| Species | Genome Information<sup>a</sup> |  |  | GR |  |  |  | NPC2 |  |  |  |
| --- | --- | --- | --- | --- | --- | --- | --- | --- | --- | --- | --- |
|  | N50 | Scaffolds | Predicted proteins | BITACORA identified sequences<sup>b</sup> | Identified in |  | Complete sequences<sup>e</sup> | BITACORA identified sequences<sup>b</sup> | Identified in |  | Complete sequences<sup>e</sup> |
|  |  |  |  |  | GFF<sup>c</sup> | Genome<sup>d</sup> |  |  | GFF<sup>c</sup> | Genome<sup>d</sup> |  |
| *C. sculpturatus* | 342,549 | 10,457 | 30,465 | 1,234 | 24 | 1,210 | 518 | 15 | 11 | 4 | 7 |
| *M. martensii* | 45,228 | 92,408 | 32,016 | 673 | 23 | 650 | 173 | 15 | 9 | 6 | 7 |
| *A. geniculata* | 20,294 | 4,986,575 | 76,238 | 137 | 2 | 135 | 54 | 20 | 14 | 6 | 13 |
| *Lo. reclusa* | 63,233 | 143,678 | 20,617 | 203 | 1 | 202 | 148 | 18 | 7 | 11 | 9 |
| *S. mimosarum* | 480,636 | 68,653 | 26,888 | 229 | 4 | 225 | 120 | 16 | 13 | 3 | 14 |
| *P. tepidariorum* | 465,572 | 31,445 | 32,186 | 819 | 52 | 767 | 383 | 20 | 19 | 1 | 17 |
| *La. hesperus* | 13,889 | 151,814 | 17,364 | 76 | 0 | 76 | 37 | 15 | 2 | 13 | 8 |
| Total chelicerata | - | - | - | 3,371 | 106 | 3,265 | 1,433 | 119 | 75 | 44 | 75 |

<sup>a</sup> Features from the focal genomes
<sup>b</sup> Total number of members of the gene family after running the two steps of BITACORA
<sup>c</sup> Total number of members of the gene family predicted in the former GFF
<sup>d</sup> Total number of new gene family members identified by BITACORA (output of Step 2)
<sup>e</sup> Number of complete sequences identified by BITACORA

The members of the NPC2 family, on the contrary, are much more conserved at the sequence level and show higher levels of gene expression in arthropods (Pelosi et al., 2014). As expected, the number of newly identified copies of this family in the seven chelicerate genomes is much lower than in the case of GRs. Even that, BITACORA was able to detect 44 new NPC2 encoding sequences, raising the total repertoire in these species to 119 (Table 1). It is worth noting, however, that a non-negligible number of these new identified genes are incomplete, likely caused either by a poor genome assembly quality (indicated as the N50 and the number of scaffolds) or a low number of annotated proteins in the input GFF requiring to predict novel gene models in BITACORA second round (only 42.5% and 63% of the uncovered GR and NPC2 proteins, respectively, were complete; Table 1), demonstrating that the performance of BITACORA depends on both the quality of input annotations and genome assemblies in addition to the specific focal gene family.

## Discussion

Gene families are one of the most abundant and dynamic components of eukaryotic genomes. Therefore, having curated genomic data is fundamental not only to carry out comprehensive comparative or functional genomics studies on gene families, but also to understand global genome architecture and biology. During the last decades, the rapid development of sequencing technologies has enabled the rapid accumulation of genome sequences of non-model organisms. Nevertheless, most of them still remain quite fragmented and only have very preliminary and incomplete automatic annotations. The proteins predicted by automatic annotation tools often contain systematic errors, such as incomplete or chimeric gene models, which are especially notable in gene families given the repetitive nature of their members. Besides, since new copies commonly arise by unequal crossing-over, they are frequently found in physically close tandem arrays of similar sequences, further complicating annotations (Clifton et al., 2017; Vieira et al., 2007).

With this in mind, we have developed a bioinformatics tool that helps researchers to access these automatic annotations, extract the information of focal gene families, curate and update gene models and identify new copies from DNA sequences. Using BITACORA, gene family annotations can be really improved using both HMM profiles and iterative searches that incorporate the new variability found in previous searches.

One of the analyses on gene families more sensitive to the quality of annotations is the estimation of the number of gene gains and losses and the associated birth and death rates. The example of the GR family in chelicerates demonstrates the importance of refining annotations using BITACORA. Indeed, using unsupervised annotations in non-model organism genomes directly to estimate turnover rates might produce very erroneous results, not only in terms of gene counts but also in calculations biased to highly expressed and/or very recent copies. Then, BITACORA can be used to reduce considerably these errors and make more accurate and robust inferences about the age/origin of the family and of its mode of evolution.

On the other hand, the curation of both existing and new identified members of a family with BITACORA might be also crucial for further analysis on their sequence evolution. The quality of multiple sequence alignments, which are used to determine orthology groups, to obtain divergence estimates or to detect the footprint of natural selection in gene family members, is strongly compromised by the presence of badly annotated copies, including chimeras and incorrectly annotated fragments. Using BITACORA we can detect these artifacts and either fix or discard them from further analyses.

Despite its proven utility, we are aware that BITACORA do not provide perfect annotations for a gene family. For this reason, we configured the pipeline output to be easily readable for genome editor tools, such as Apollo, which facilitate researchers to improve gene models. Fig. 3 show an example of the annotation tracks generated by BITACORA (BED files) for a member of the candidate carrier protein (*Ccp*: Vizueta et al., 2017) in the genome draft of the spider *Dysdera silvatica* (unpublished data). The automatic annotation using MAKER2 (track GFF3 Dsil) generated an incomplete gene model (with three missing putative exons) that could be easily improved given its identification with BITACORA and the generated output.

**Fig. 2.**
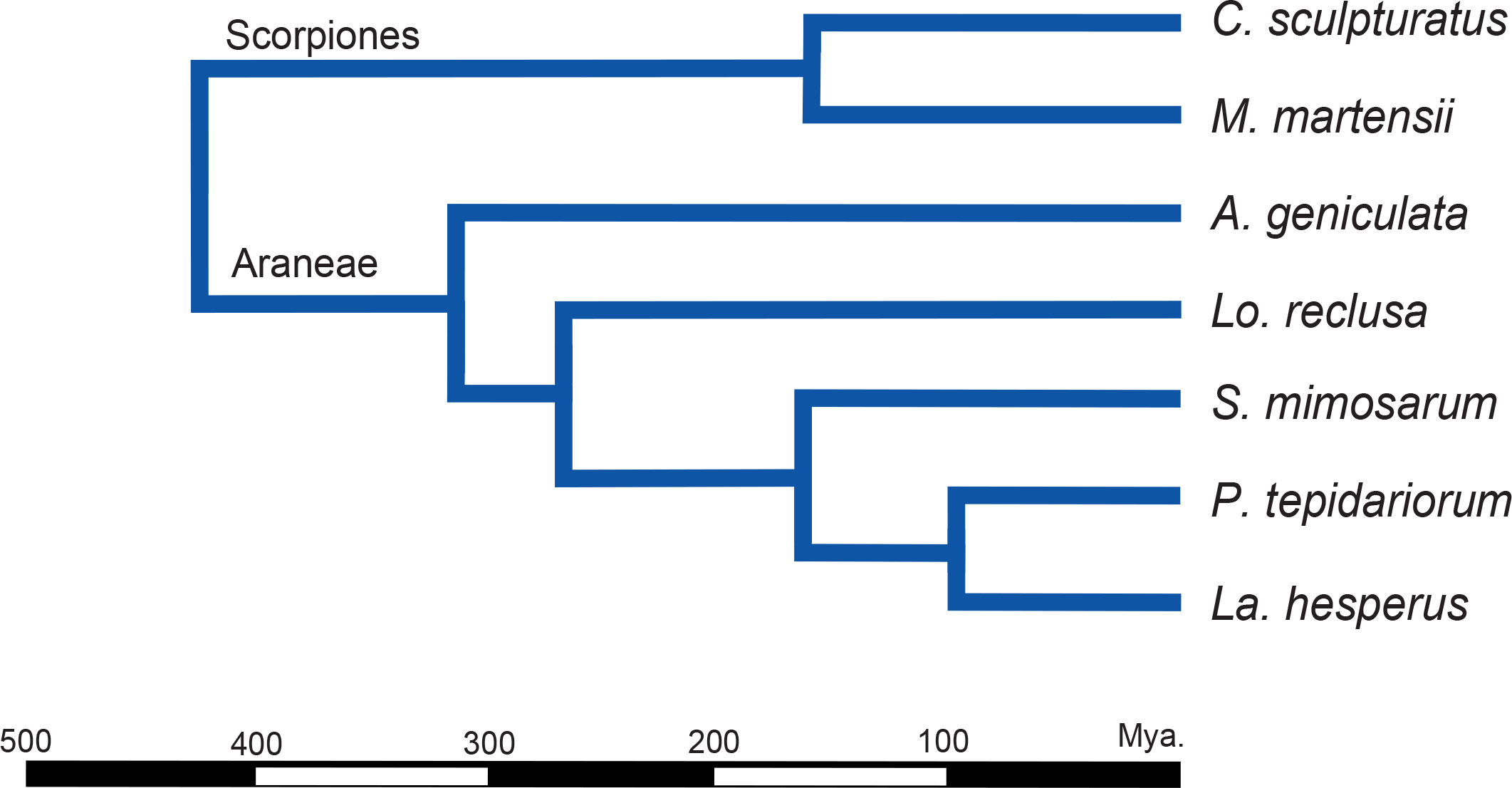
Phylogenetic relationships among the seven chelicerate species surveyed for the GR and the NPC2 families.

**Fig. 3.**
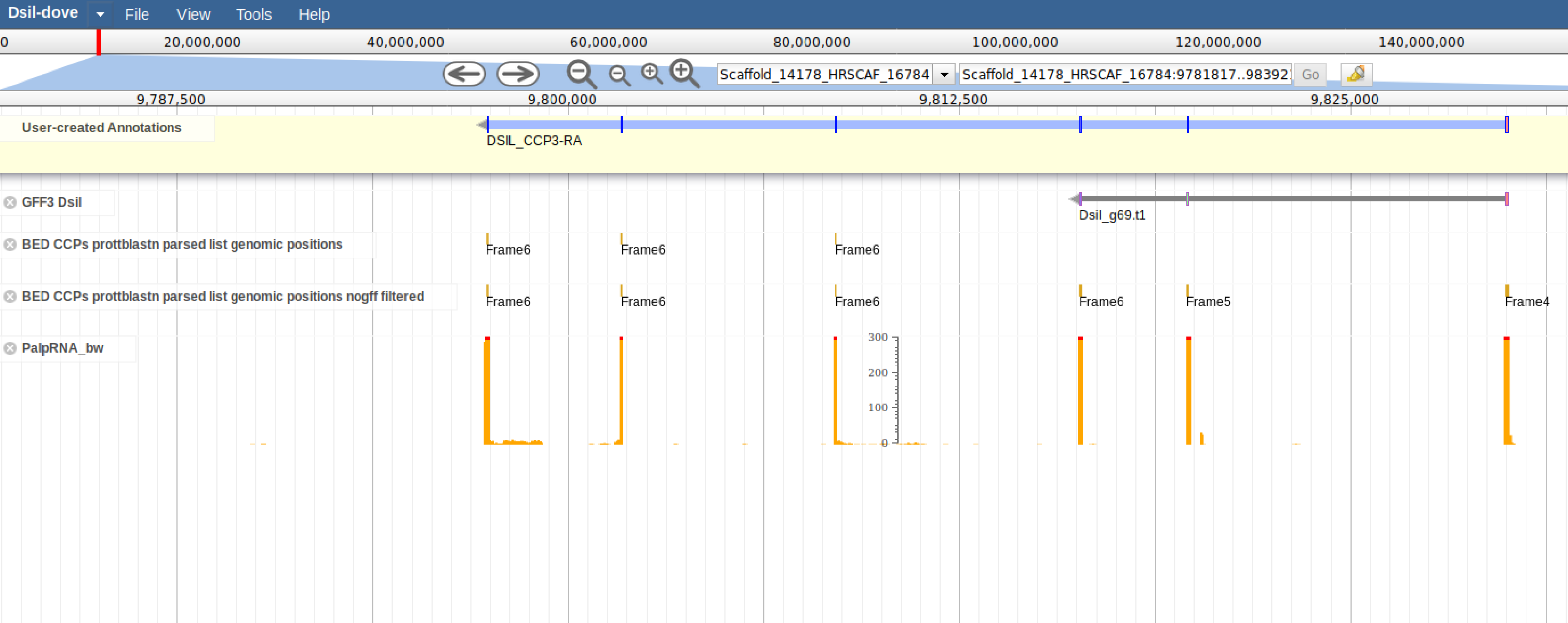
Visualization in Apollo genome editor of the BITACORA output with the annotation features of a candidate carrier protein (*Ccp*) gene (Dsil_g69.t1 in *D. silvatica*). The Dsil GFF3 track shows the original GFF file obtained by BRAKER in the focal genome. The two BED tracks shows the output files generated in BITACORA showing all putative exons identified in a region not annotated in the original GFF and the all six exons identified by BITACORA. The RNA track (PalpRNA_bw) shows the genomic regions with mapped reads from the sequencing of aspider palp RNA-seq library. The final gene model is shown in the User-created Annotations track.

## Conclusion

Genome annotation, especially in non-model organisms, is still a bottleneck for evolutionary and functional genomic analyses. To assists this task, we developed a comprehensive pipeline that facilitates the curation of existing models and the identification of new gene family copies in genome assemblies with available annotation features and/or genomic sequences. The output of BITACORA can be used as a baseline for manual annotation in genomic annotation editors, used as evidence in automatic annotation tools to improve gene family model predictions, or to directly perform downstream analysis. Future directions should include the implementation in our pipeline of an automatic annotation tool to directly predict new gene models in DNA sequences and its integration as a part of genome annotation editors to facilitate gene family annotation in collaborative genome projects.

## Acknowledgements

We would like to thank Paula Escuer and Vadim Pisarenco for helpful discussions. This work was supported by the Ministerio de Economía y Competitividad of Spain (CGL2013-45211, CGL2016-75255) and the Comissió Interdepartamental de Recerca I Innovació Tecnològica of Catalonia, Spain (2017SGR1287). J.V. was supported by a FPI grant (Ministerio de Economía y Competitividad of Spain, BES-2014-068437).

## Author contributions

J.V., A.S.-G and J.R. conceived the work. J.V. wrote the scripts, did the analyses and wrote the first version of the manuscript. All authors checked and confirmed the final version of the manuscript.

## Data accessibility

BITACORA is available from http://www.ub.edu/softevol/bitacora, and https://github.com/molevol-ub/bitacora

## Supplementary Material

BITACORA Documentation

## References

Altschul, S. (1997). Gapped BLAST and PSI-BLAST: a new generation of protein database search programs. Nucleic Acids Research, 25(17), 3389–3402. doi:10.1093/nar/25.17.3389

Clifton, B. D., Librado, P., Yeh, S.-D., Solares, E. S., Real, D. A., Jayasekera, S. U., … Ranz, J. M. (2017). Rapid Functional and Sequence Differentiation of a Tandemly Repeated Species-Specific Multigene Family in *Drosophila*. Molecular Biology and Evolution, 34(1), 51–65. doi:10.1093/molbev/msw212

Dominguez Del Angel, V., Hjerde, E., Sterck, L., Capella-Gutierrez, S., Notredame, C., Vinnere Pettersson, O., … Lantz, H. (2018). Ten steps to get started in Genome Assembly and Annotation. F1000Research, 7, ELIXIR–148. doi:10.12688/f1000research.13598.1

Eddy, S. R. (2011). Accelerated Profile HMM Searches. PLoS Computational Biology, 7(10), e1002195. doi:10.1371/journal.pcbi.1002195

Finn, R. D., Bateman, A., Clements, J., Coggill, P., Eberhardt, R. Y., Eddy, S. R., … Punta, M. (2014). Pfam: the protein families database. Nucleic Acids Research, 42(Database issue), D222–D230. doi:10.1093/nar/gkt1223

Gremme, G., Brendel, V., Sparks, M. E., & Kurtz, S. (2005). Engineering a software tool for gene structure prediction in higher organisms. Information and Software Technology, 47(15), 965–978. doi:10.1016/J.INFSOF.2005.09.005

Hoff, K. J., Lange, S., Lomsadze, A., Borodovsky, M., & Stanke, M. (2016). BRAKER1: Unsupervised RNA-Seq-Based Genome Annotation with GeneMark-ET and AUGUSTUS. Bioinformatics, 32(5), 767–769. doi:10.1093/bioinformatics/btv661

Holt, C., & Yandell, M. (2011). MAKER2: an annotation pipeline and genome-database management tool for second-generation genome projects. BMC Bioinformatics, 12(1), 491. doi:10.1186/1471-2105-12-491

Joseph, R. M., & Carlson, J. R. (2015). *Drosophila* Chemoreceptors: A Molecular Interface Between the Chemical World and the Brain. Trends in Genetics□: TIG, 31(12), 683–695. doi:10.1016/j.tig.2015.09.005

Keller, O., Kollmar, M., Stanke, M., & Waack, S. (2011). A novel hybrid gene prediction method employing protein multiple sequence alignments. Bioinformatics, 27(6), 757–763. doi:10.1093/bioinformatics/btr010

Korf, I. (2004). Gene finding in novel genomes. BMC Bioinformatics, 5, 59. doi:10.1186/1471-2105-5-59

Lee, E., Helt, G. A., Reese, J. T., Munoz-Torres, M. C., Childers, C. P., Buels, R. M., … Lewis, S. E. (2013). Web Apollo: a web-based genomic annotation editing platform. Genome Biology, 14(8), R93. doi:10.1186/gb-2013-14-8-r93

Lomsadze, A., Burns, P. D., & Borodovsky, M. (2014). Integration of mapped RNA-Seq reads into automatic training of eukaryotic gene finding algorithm. Nucleic Acids Research, 42(15), e119–e119. doi:10.1093/nar/gku557

Pelosi, P., Iovinella, I., Felicioli, A., & Dani, F. R. (2014). Soluble proteins of chemical communication: an overview across arthropods. Frontiers in Physiology, 5(August), 320. doi:10.3389/fphys.2014.00320

Robertson, H. M. (2015). The Insect Chemoreceptor Superfamily Is Ancient in Animals. Chemical Senses, 40(9), 609–614. doi:10.1093/chemse/bjv046

Slater, G. S. C., & Birney, E. (2005). Automated generation of heuristics for biological sequence comparison. BMC Bioinformatics, 6, 31. doi:10.1186/1471-2105-6-31

Stanke, M., & Waack, S. (2003). Gene prediction with a hidden Markov model and a new intron submodel. Bioinformatics, 19(Suppl 2), ii215–ii225. doi:10.1093/bioinformatics/btg1080

Stanke, Mario, Diekhans, M., Baertsch, R., & Haussler, D. (2008). Using native and syntenically mapped cDNA alignments to improve de novo gene finding. Bioinformatics, 24(5), 637–644. doi:10.1093/bioinformatics/btn013

Vieira, F. G., Sánchez-Gracia, A., & Rozas, J. (2007). Comparative genomic analysis of the odorant-binding protein family in 12 *Drosophila* genomes: purifying selection and birth-and-death evolution. Genome Biology, 8(11), R235. doi:10.1186/gb-2007-8-11-r235

Vizueta, J., Frías-López, C., Macías-Hernández, N., Arnedo, M. A., Sánchez-Gracia, A., & Rozas, J. (2017). Evolution of chemosensory gene families in arthropods: Insight from the first inclusive comparative transcriptome analysis across spider appendages. Genome Biology and Evolution, 9(1), 178–196. doi:10.1093/gbe/evw296

Vizueta, J., Rozas, J., & Sánchez-Gracia, A. (2018). Comparative Genomics Reveals Thousands of Novel Chemosensory Genes and Massive Changes in Chemoreceptor Repertories across Chelicerates. Genome Biology and Evolution, 10(5), 1221–1236. doi:10.1093/gbe/evy081

Yandell, M., & Ence, D. (2012). A beginner’s guide to eukaryotic genome annotation. Nature Reviews Genetics, 13(5), 329–342. doi:10.1038/nrg3174

Yohe, L. R., Davies, K. T. J., Simmons, N. B., Sears, K. E., Dumont, E. R., Rossiter, S. J., & Dávalos, L. M. (2019). Evaluating the performance of targeted sequence capture, RNA-Seq, and degenerate-primer PCR cloning for sequencing the largest mammalian multigene family. Molecular Ecology Resources. doi:10.1111/1755-0998.13093

Zhang, Y., Zheng, Y., Li, D., & Fan, Y. (2014). Transcriptomics and identification of the chemoreceptor superfamily of the pupal parasitoid of the oriental fruit fly, *Spalangia endius* Walker (Hymenoptera: Pteromalidae). PloS One, 9(2), e87800. doi:10.1371/journal.pone.0087800

