## Supplementary Material for "BITACORA: A comprehensive tool for the identification and annotation of gene families in genome assemblies"

### BITACORA:

Departament de Genètica, Microbiologia i Estadística
Institut de Recerca de la Biodiversitat (IRBio)

**Universitat de Barcelona**

http://www.ub.es/softevol/bitacora

September 17th, 2019

UNIVERSITAT DE BARCELONA

EG&B Evolutionary Genomics & Bioinformatics

#### - Overview

Genome annotation is a critical bottleneck in genomic research, especially for the rigorous and comprehensive study of gene families. Despite current progress in automatic annotation, the tools developed for this task often generate absent or inaccurate gene models, such as fused or chimeric genes, which require an extra substantial effort to be correctly annotated. Here we present BITACORA, a bioinformatic tool that integrates sequence similarity search algorithms and Perl scripts to facilitate the curation of annotation errors, and the de novo identification of new family copies in genome sequences. The pipeline generates general feature format (GFF) files with both curated and novel gene models and FASTA files with the predicted proteins. The output of BITACORA can be easily integrated in genomic annotation editors, greatly facilitating subsequent semi-automatic annotation and downstream analyses.

**Authors**

| | |
|---|---|
| Joel Vizueta | |
| Alejandro Sánchez-Gracia | |
| Julio Rozas | |


##### BITACORA Publication

Vizueta, J., Sánchez-Gracia, A. and Rozas, J. 2019. BITACORA: A comprehensive tool for the identification and annotation of gene families in genome assemblies. *Submitted*.

##### BITACORA Web Site

www.ub.edu/softevol/bitacora

### 0 Workflow & Contents

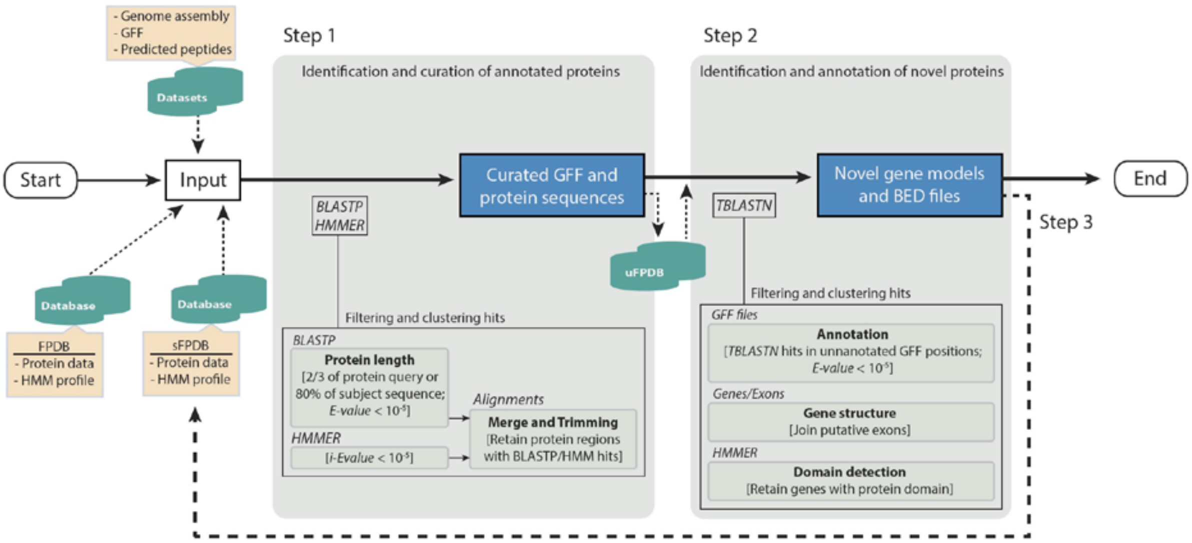


Figure. Workflow showing the basic steps used in BITACORA


**1.** Installation

**2.** Prerequisites

**3.** Computational Requirements

**4.** Usage modes

- **4.1.** Full mode
- **4.2.** Protein mode
- **4.3.** Genome mode

**5.** Parameters

**6.** Running BITACORA

**7.** Output

**8.** Example

**9.** Citation

**10.** Troubleshooting

### 1 Installation

BITACORA is distributed as a multiplatform shell script (runBITACORA.sh) that calls several other perl scripts, which include all functions responsible of performing all pipeline tasks. Hence, it does not require any installation or compilation step.

You can download all package contents from GitHub: https://github.com/molevol-ub/bitacora

To run the pipeline edit the master script runBITACORA.sh variables described in Prerequisites, Data, and Parameters.

#### 2 Prerequisites

- **BLAST**: Download blast executables from:
ftp://ftp.ncbi.nlm.nih.gov/blast/executables/blast+/LATEST/

- **HMMER**: The easiest way to install HMMER in your system is to type one of the following commands in your terminal:

```
% brew install hmmer                  # OS/X, HomeBrew
% port install hmmer                  # OS/X, MacPorts
% apt install hmmer                   # Linux (Ubuntu, Debian...)
% dnf install hmmer                   # Linux (Fedora)
% yum install hmmer                   # Linux (older Fedora)
% conda install -c bioconda hmmer     # Anaconda
```

Or compile HMMER binaries from the source code: http://hmmer.org/

- **Perl**: Perl is installed by default in most operating systems. See https://learn.perl.org/installing/ for installation instructions.

HMMER and BLAST binaries require to be added to the PATH environment variable. Specify the correct path to bin folders in the master script `runBITACORA.sh`, if necessary.

```
$ export PATH=$PATH:/path/to/blast/bin
$ export PATH=$PATH:/path/to/hmmer/bin
```

### 3 Computational requirements

BITACORA have been tested in UNIX-based platforms (both in Mac OS and Linux operating systems). Multiple threading can be set in blast searches, which is the most time-consuming step, by editing the option THREADS in `runBITACORA.sh`

For a typical good quality genome (~2Gb in size and ~10,000 scaffolds) and a standard modern PC (16Gb RAM), a full run of BITACORA is completed in less than 24h. This running time, however, will depend on the size of the gene family or the group of genes surveyed in a particular analysis. For gene families of 10 to 100 members, BITACORA spends from minutes to a couple of hours.

In case of larger or very fragmented genomes, BITACORA should be used in a computer cluster or workstation given the increase of RAM memory and time required.

### 4 Usage modes

#### 4.1. Full mode

BITACORA has been initially designed to work with genome sequences and protein annotations (full mode). However, the pipeline can also be used either with only protein or only genomic sequences (protein and genome modes, respectively). These last modes are explained in next subsections.

**Preparing the data**: The input files (in plain text) required by BITACORA to run a full analysis are (update the complete path to these files in the master script `runBITACORA.sh`):

**I**. File with genomic sequences in FASTA format

**II**. File with structural annotations in GFF3 format. [*NOTE: mRNA* or *transcript*, and *CDS* are mandatory fields].

-------------------- GFF3 example

```
lg1_ord1_scaf1770	AUGUSTUS	gene	13591	13902	0.57	+	.	ID=g1;
lg1_ord1_scaf1770	AUGUSTUS	mRNA	13591	13902	0.57	+	.	ID=g1.t1;Parent=g1;
lg1_ord1_scaf1770	AUGUSTUS	start_codon	13591	13593	.	+	0	Parent=g1.t1;
lg1_ord1_scaf1770	AUGUSTUS	CDS	13591	13902	0.57	+	0	ID=g1.t1.CDS1;Parent=g1.t1
lg1_ord1_scaf1770	AUGUSTUS	exon	13591	13902	.	+	.	ID=g1.t1.exon1;Parent=g1.t1;
lg1_ord1_scaf1770	AUGUSTUS	stop_codon	13900	13902	.	+	0	Parent=g1.t1;
```

--------------------

BITACORA also accepts other GFF formats, such as Ensembl GFF3 or GTF. [NOTE: GFF formatted files from NCBI can cause errors when processing the data, use the supplied script "`reformat_ncbi_gff.pl`" (located in the folder `/Scripts/Tools`) to make the file parsable by BITACORA]. See Troubleshooting in case of getting errors while parsing your GFF.

-------------------- Ensembl GFF3 example

```
AFFK01002511	EnsemblGenomes	gene	761	1018	.	-	.	ID=gene:SMAR013822;assembly_name=Smar1;biotype=protein_coding;logic_name=ensemblgenomes;versio
AFFK01002511	EnsemblGenomes	transcript	761	1018	.	-	.	ID=transcript:SMAR013822-RA;Parent=gene:SMAR013822;assembly_name=Smar1;biotype=protein
AFFK01002511	EnsemblGenomes	CDS	761	811	.	-	0	Parent=transcript:SMAR013822-RA;assembly_name=Smar1
AFFK01002511	EnsemblGenomes	exon	761	811	.	-	.	Parent=transcript:SMAR013822-RA;Name=SMAR013822-RA-E2;assembly_name=Smar1;constitutive=1;ensem
AFFK01002511	EnsemblGenomes	CDS	887	1018	.	-	0	Parent=transcript:SMAR013822-RA;assembly_name=Smar1
AFFK01002511	EnsemblGenomes	exon	887	1018	.	-	.	Parent=transcript:SMAR013822-RA;Name=SMAR013822-RA-E1;assembly_name=Smar1;constitutive=1;ensem
```

--------------------

**III**. File with predicted proteins in FASTA format. BITACORA requires identical IDs for proteins and their corresponding mRNAs or transcripts IDs in the GFF3. [NOTE: we recommend using genes but not isoforms in BITACORA; isoforms can be removed or properly annotated after BITACORA analysis]

**IV**. Specific folder with files containing the query protein databases (`YOURFPDB_db.fasta`) and HMM profiles (`YOURFPDB_db.hmm`) in FASTA and hmm format, respectively, where the "YOURFPDB" label is your specific data file name. The

addition of "_db" to the database name with its proper extension, `fasta` or `hmm`, is mandatory.
BITACORA requires one protein database and profile per surveyed gene family (or gene group). See `Example/DB` files for an example of searching for two different gene families in BITACORA: OR, Odorant Receptors; and CD36-SNMP.

[NOTE: profiles covering only partially the proteins of interest are not recommended]

<u>Notes on HMM profiles:</u>

HMM profiles are found in InterPro or PFAM databases associated to known protein domains. If you don't know if your protein contains any described domain, you can search in InterPro (http://www.ebi.ac.uk/interpro/) using the protein sequence of one of your queries to identify domains.

For example, for the chemosensory proteins (CSPs) in insects, you can download the HMM profile from pfam (Curation & model PFAM submenu):

http://pfam.xfam.org/family/PF03392#tabview=tab6

In the case of searching for proteins with not described protein domains, or with domains not covering most of the protein sequence, it should be performed an alignment of the query proteins to create a specific HMM profile.

Example of building a protein profile (it requires an aligner, here we use mafft as example):

```
$ mafft --auto FPDB_db.fasta > FPDB_db.aln
$ hmmbuild FPDB_db.hmm FPDB_db.aln
```

<u>Notes on the importance of selecting a confident curated database:</u>

The proteins included in the database to be used as query (FPDB) in the protein search is really important; indeed, the inclusion of unrelated or bad annotated proteins could lead to the identification and annotation of proteins unrelated to the focal gene family and can inflate the number of sequences identified.

On the other hand, if possible, we recommend to include proteins from phylogenetically-close species to increase the power of identifying proteins, particularly in fast-evolving and divergent gene families. If your organism of interest does not have an annotated genome of a close related species, we suggest to perform a second BITACORA round (step 3 described in the manuscript), including in the query database (sFPDB) the sequences identified in the first round, along with a new HMM profile build with these sequences. This step may facilitate the identification of previously undetected related divergent sequences.

#### 4.2. Protein mode

BITACORA can also run with a set of proteins (i.e. predicted proteins from transcriptomic data; script `runBITACORA_protein_mode.sh`) by using the input files described in points **III** and **IV** of the section **4.1**.

Under this mode, BITACORA identifies, curates when necessary, and report all members of the surveyed family among the predicted proteins. The original protein sequences (not being curated) are also reported (located in `Intermediate_Files` if cleaning output is active).

#### 4.3. Genome mode

BITACORA can also run with raw genome sequences (i.e., not annotated genomes; script `runBITACORA_genome_mode.sh`), by using the input files described in points **I** and **IV** of the section **4.1**.

Under this mode, BITACORA identifies *de novo* all members of the surveyed family and returns a BED file with gene coordinates of the detected exons, a FASTA file with predicted proteins from these exons and a GFF3 file with the corresponding structural annotations.

[NOTE: The gene models generated under this mode are only semi-automatic predictions and require further manual annotation, i.e. using genomic annotation editors, such as Apollo. The output file of the genome mode can also be used as protein evidence in automatic annotators as MAKER2 or BRAKER1 (see output section)]

### 5 Parameters

- The option CLEAN can be used to create the `Intermediate_files` directory where all intermediate files will be stored (see output section).

CLEAN=T #T=true, F=false

- BLAST and HMMER hits are filtered with a default cut-off E-value of 10e-5 (in addition to an internal parameter for filtering the length covered by the alignment).

E-value can be modified in the master script runBITACORA.sh:

EVALUE=10e-5

- Number of threads to be used in blast searches, default is 1.

THREADS=1

- BITACORA uses by default a value of 15 kb to join putative exons from separate but contiguous (and in the same scaffold) genome hits.

This value can be modified in the master script runBITACORA.sh:

MAXINTRON=15000

Notes on the parameter MAXINTRON:

**Estimating the intron length distribution in your genome**:

MAXINTRON is a critical parameter affecting the quality of the gene models built after joining *de novo* identified exons after BLASTN search (see BITACORA article). BITACORA is distributed with a script (`get_intron_size_fromgff.pl`) to compute some summary statistics, such as the mean, median, and the 95% and 99% upper limits of the intron length distribution, of an input GFF, which can contain all genes from genome or only the genes identified for a particular gene family (i.e. GFF generated in BITACORA output).

Note that a very high value could join exons from different genes, generating a putative chimeric gene. On the other hand, a very low value could not join exons from the same gene. Therefore, it is very important to set a MAXINTRON biological realistic value, which could vary across species or assemblies. As default, BITACORA uses a conservative high value, as a compromise between ensuring the joining of all exons from a same gene, and avoiding the generation of erroneous gene fusions. In any case, a large value of MAXINTRON parameter prevents the annotation of fragmented genes but can generate gene models with multiple gene fusions. Putative gene fusions (proteins with two or more domains predicted by BITACORA) are tagged with the label "Xdom" at the end of the protein name in the output file, being X the number of putative genes (detected domains).

The number of putative fussed genes identified as new proteins in not annotated regions of the genome can be obtained using the following command in the terminal:

```
$ grep '>.*dom' DB/DB_genomic_and_annotated_proteins_trimmed.fasta
```

### 6 Running BITACORA

After preparing the data as indicated in steps 5 (Usage) and 6 (Parameters), you can execute BITACORA with the following command:

```
$ bash runBITACORA.sh
```

### 7 Output

BITACORA creates an output folder for each query database, and three files with the number of proteins identified in each step, including a summary table. For the genome and protein modes, only one summary table will be reported with the number of identified genes.

In each folder, there are the following **main files** (considering you chose to clean output directory. If not, all files will be found in the same output folder):

- `YOURFPDB_genomic_and_annotated_genes_trimmed.gff3`: GFF3 file with information of all identified protein curated models both in already annotated proteins and unannotated genomic sequences.

- `YOURFPDB_genomic_and_annotated_proteins_trimmed.fasta`: A fasta file containing the protein sequences from the above gene models.

**Non-redundant data**: Relevant information excluding identical proteins, or those considered as artefactual false positives (i.e. duplicated scaffolds, isoforms…).

- `YOURFPDB_genomic_and_annotated_genes_trimmed_nr.gff3`: GFF3 file containing all identified non-redundant protein curated models both in already annotated proteins and unannotated genomic sequences.

- `YOURFPDB_genomic_and_annotated_proteins_trimmed.fasta`: A fasta file containing the non-redundant protein sequences from the above gene models.

**BED files** with non-redundant merged blast hits in genome sequence:

- `YOURFPDBtblastn_parsed_list_genomic_positions.bed`: BED file with only merged blast alignments in non-annotated regions.

- `YOURFPDBtblastn_parsed_list_genomic_positions_nogff_filtered.bed`: BED file with merged blast alignment in all genomic regions.

In addition, BITACORA generates the following **Intermediate files** (located into Intermediate_files folder created in cleaning, if active):

- `YOURFPDB_annot_genes.gff3` and `YOURFPDB_proteins.fasta`: GFF3 and fasta file containing the original untrimmed models for the identified proteins.

- `YOURFPDB_annot_genes_trimmed.gff3` and `YOURFPDB_proteins_trimmed.fasta`: GFF3 and fasta containing only the curated model for the identified annotated proteins (trimming exons if not aligned to query FPDB sequences or split putative fused genes)

- YOURFPDB_genomic_genes.gff3: GFF3 containing novel identified proteins in genomic sequences.

- YOURFPDB_genomic_genes_trimmed.gff3: GFF3 containing novel identified proteins in genomic sequences curated by the positions identified in the HMM profile.

- YOURFPDBgfftrimmed.cds.fasta and YOURFPDBgfftrimmed.pepfasta: Files containing CDS and protein sequences translated directly from YOURFPDB_annot_genes_trimmed.gff3

- YOURFPDBgffgenomictrimmed.cds.fasta and YOURFPDBgffgenomictrimmed.pep.fasta: Files containing CDS and protein sequences translated directly from YOURFPDB_genomic_genes_trimmed.gff3

- hmmer folder containing the output of HMMER searches against the annotated proteins and novel proteins identified in the genome

- YOURFPDB_blastp.outfmt6: BLASTP output of the search of the query FPDB against the annotated proteins

- YOURFPDB_tblastn.outfmt6: TBLASTN output of the search of the query FPDB against the genomic sequence

- YOURFPDB_blastp_parsed_list.txt; YOURFPDB_hmmer_parsed_list.txt; YOURFPDB_allsearches_list.txt; YOURFPDB_combinedsearches_list.txt: Parsed files combining all hits and extending the hit positions from BLASTP and HMMER outputs

- YOURFPDB_tblastn_parsed_list_genomic_positions.txt (and _notgff_filtered): File containing the positions identified after parsing the TBLASTN search.

- YOURFPDB_prots_VsGFF_badannot_list.txt and _goodannot_list.txt: Debugging files: These files are for checking that the identified proteins and the protein models in the GFF3 codify the same protein. If the file badannot_list.txt contains some identifier, it means that the GFF3 annotation is incorrect pointing to a bad annotation in the original GFF3. Please, try to translate the CDS for that protein into the 3 reading frames and check if the 2nd or 3rd frame codify for the protein in question stored in "YOURFPDB_genomic_and_annotated_proteins_trimmed_nr.fasta". If correct, modify the GFF3 by adding 1 or 2 nucleotide position in the start of the GFF3 (take into account if it is transcribed from forward or reverse strand). If negative, please report the error via GitHub.

- YOURFPDB_genomic_genes_proteins.fasta: It contains all merged exons from putative novel proteins identified in the genome before filtering those without the protein domain identified with HMMER. This file could be useful in case of using an HMM profile not trained with your sequences which cannot detect divergent sequences, such as remote homologs.

- YOURFPDB_genomic_exon_proteins.fasta: contains the exon sequences joined into genes in the aforementioned file.

- Additional generated files are stored for pipeline debugging and controls.

<u>Notes on BITACORA output:</u>

The obtained proteins could be used for further prospective analyses or to facilitate a more curated annotation using genome annotation editors or, in the case of having a high number of not annotated proteins in the GFF, BITACORA output sequences could also be used as evidence to improve the annotation of automatic annotators as MAKER2 or BRAKER1. However, a first validation of the obtained proteins should be performed, more specifically in those obtained newly from genome (taking into account the parameter used to join putative exons, to split putative joined genes or join exons from the same gene). In addition, these proteins obtained and assembled from genomic regions are illustrative, but more putative genes (true negatives) could be obtained from the TBLASTN BED file positions discarded for not being identified with the protein domain (i.e. alignments containing introns between two proximal exons could lead not to identify the domain in the protein).

Such validation to identify putative erroneously assigned proteins (mainly caused by the inclusion of contaminant sequences in the query database) could consist in aligning all proteins and checking the MSA, constructing the phylogeny of the gene with related species or the gene family; doing a reduced blast with NCBI-nr database or obtaining structural particularities of the proteins (i.e. characterizing protein domains as transmembrane domains, signal peptides...). See our manuscript Vizueta et al. (2018) for an example of such analyses.

In particular, BITACORA full and genome mode is also designed to facilitate the gene annotation in editors as Apollo. For that, the use of the following files would be useful (see an example in Documentation/example_Apollo.png):

- Original GFF3
- Final GFF3 with curated models for the annotated proteins
- BED file from TBLASTN search

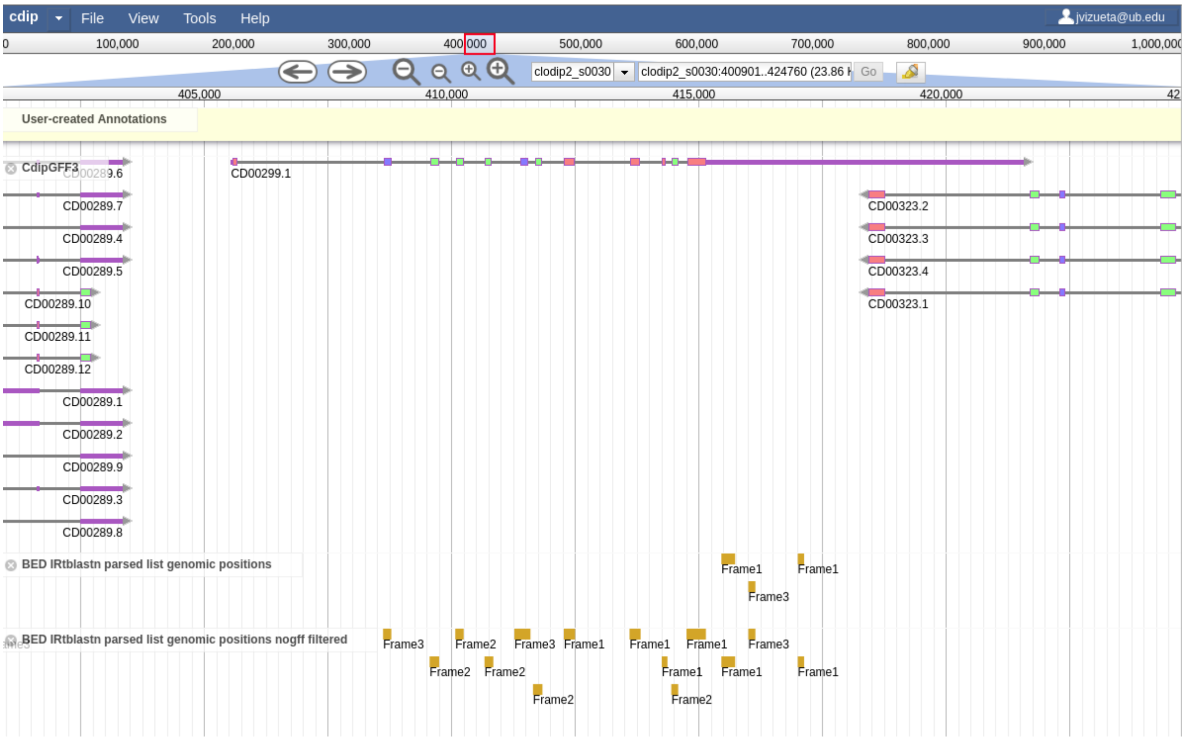

If there are sequences containing stop codons, codified as "X", it could be artefactual from TBLASTN hits if they are in the beginning or end of an exon or, otherwise, those genes are probably pseudogenes.

Nonetheless, again, new proteins identified from unannotated genomic regions should be properly annotated using genome browser annotation tools such as Apollo, or could be used as evidence to improve the annotation of automatic annotators as MAKER2 or BRAKER1. We estimate an approximate number of them which could be used for prospective analyses.

### 8 Example

An example to run BITACORA can be found in `Example` folder. First, unzip the `Example_files.zip` file to obtain the necessary files for BITACORA. In this example, two chemosensory-related gene families in insects: Odorant receptors (ORs), and the CD36-SNMP gene family; will be searched in the chromosome 2R of *Drosophila melanogaster*. The GFF3 and protein files are modified from original annotations, deleting some gene models, to allow that BITACORA can identify novel not-annotated genes.

To run the example, edit the master script `runBITACORA.sh` to add the path to BLAST and HMMER binaries and run the script. It will take around 1 minute with 2 threads.

```
$ bash runBITACORA.sh
```

### 9 Citation

Joel Vizueta, Alejandro Sánchez-Gracia, and Julio Rozas. 2019. BITACORA: A comprehensive tool for the identification and annotation of gene families in genome assemblies. *Submitted*.

Moreover, you can also cite the following article where we describe the protein annotation procedure:

Joel Vizueta, Julio Rozas, Alejandro Sánchez-Gracia; Comparative Genomics Reveals Thousands of Novel Chemosensory Genes and Massive Changes in Chemoreceptor Repertories across Chelicerates, Genome Biology and Evolution, Volume 10, Issue 5, 1 May 2018, Pages 1221–1236, https://doi.org/10.1093/gbe/evy081

### 10 Troubleshooting

When BITACORA detects any error related to input data, it stops and prints the description of the error. Please check the error and your data.

If you are getting errors related to parsing the GFF file, take into account that BITACORA expects proteins ID to be as ID in mRNA rows from GFF3.

In case of protein ID and mRNA ID causing error as they are not the named equally, first, you can use the script located in `Scripts/Tools/get_proteins_notfound_ingff.pl` to check which proteins are not found in the GFF3 file, as detailed in the Error message. You could use only those proteins found in the GFF3 in BITACORA.

If all proteins are named differently in the GFF3, you can obtain a protein file from the GFF3 using the script `Scripts/gff2fasta_v3.pl` and use that protein file as input to BITACORA.

You could also modify the perl module `Readgff.pm` to allow BITACORA to read your data. Otherwise, modify the GFF, preferably, as GFF3 format.

If you cannot solve the error, create an issue in Github specifying the error and all details as possible.
